# A new method for counting reproductive structures in digitized herbarium specimens using Mask R-CNN

**DOI:** 10.1101/2020.06.25.169888

**Authors:** Charles Davis, Julien Champ, Daniel S. Park, Ian Breckheimer, Goia M. Lyra, Junxi Xie, Alexis Joly, Dharmesh Tarapore, Aaron M. Ellison, Pierre Bonnet

## Abstract

Phenology–the timing of life-history events–is a key trait for understanding responses of organisms to climate. The digitization and online mobilization of herbarium specimens is rapidly advancing our understanding of plant phenological response to climate and climatic change. The current practice of manually harvesting data from individual specimens, however, greatly restricts our ability to scale-up data collection. Recent investigations have demonstrated that machine-learning approaches can facilitate this effort. However, present attempts have focused largely on simplistic binary coding of reproductive phenology (e.g., presence/absence of flowers). Here, we use crowd-sourced phenological data of buds, flowers, and fruits from *>* 3000 specimens of six common wildflower species of the eastern United States (*Anemone canadensis* L., *A. hepatica* L., *A. quinquefolia* L., *Trillium erectum* L., *T. grandiflorum* (Michx.) Salisb., and *T. undulatum* Wild.) to train models using Mask R-CNN to segment and count phenological features. A single global model was able to automate the binary coding of each of the three reproductive stages with *>* 87% accuracy. We also successfully estimated the relative abundance of each reproductive structure on a specimen with ≥ 90% accuracy. Precise counting of features was also successful, but accuracy varied with phenological stage and taxon. Specifically, counting flowers was significantly less accurate than buds or fruits likely due to their morphological variability on pressed specimens. Moreover, our Mask R-CNN model provided more reliable data than non-expert crowd-sourcers but not botanical experts, highlighting the importance of high-quality human training data. Finally, we also demonstrated the transferability of our model to automated phenophase detection and counting of the three *Trillium* species, which have large and conspicuously-shaped reproductive organs. These results highlight the promise of our two-phase crowd-sourcing and machine-learning pipeline to segment and count reproductive features of herbarium specimens, thus providing high-quality data with which to investigate plant response to ongoing climatic change.

## 2 Introduction

Climate change is a potent selective force that is shifting the geographic ranges of genotypes, altering population dynamics of individual species, and reorganizing entire assemblages in all environments. A key functional trait in this regard is phenology: the timing of life-history events, such as the onset of flowering or migration. The use of museum specimens has invigorated and enriched the investigation of phenological responses to climatic change, and is one of several research directions that has brought a renewed sense of purpose and timeliness to natural history collections (Meineke et al., 2018, 2019; Willis et al., 2017; Davis et al., 2015; Hedrick et al., 2020). Herbarium specimens greatly expand the historical depth, spatial scale, and species diversity of phenological observations relative to those available from field observations (Wolkovich et al., 2014). In many cases, herbarium specimens provide the only means of assessing phenological responses to climatic changes occurring over decades to centuries (Davis et al., 2015). However, a great challenge in using these specimens is accessing and rapidly assessing phenological state(s) of the world’s estimated 393 million herbarium specimens (Thiers, 2017; Sweeney et al., 2018).

The ongoing digitization and online mobilization of herbarium specimens has facilitated their broad access with significant economies of scale (Nelson and Ellis, 2019; Sweeney et al., 2018; Hedrick et al., 2020) and accelerated advances in scientific investigations, including phenological assessment efforts that were underway prior to mass digitization (Davis et al., 2015; Miller-Rushing et al., 2006; Primack et al., 2004). Digitization 2.0 (*sensu* Hedrick et al., 2020) has also sparked the integration and development of new scholarly disciplines and lines of inquiry not possible previously. Whereas Digitization 1.0 refers to the generation of digitized products from physical specimens, Digitization 2.0 is the use of natural history collections to answer scientific questions using only their digitized representation, rather than the physical specimen itself.

In recent years, scientists have used these digitized herbarium specimens in novel ways (*e*.*g*., Meineke et al., 2018, 2019; Hedrick et al., 2020) and greatly increased the pace at which key phenological trait data can be harvested from tens of thousands of specimens. *CrowdCurio*–*Thoreau’s Field Notes* (Willis et al., 2017) was one of the first attempts to move beyond the standard practice of coding phenology of herbarium specimens using binary (presence/absence) coding (*e*.*g*., specimen A has flowers, specimen B has fruits: Miller-Rushing et al., 2006; Primack et al., 2004). Many of these efforts have also focused largely on flowering, ignoring other key phenophases. Rather, users of *CrowdCurio* use a crowd-sourcing pipeline to score and quantify all phenophase features–bud, flowers, and fruits–for each specimen processed. This pipeline has facilitated the first development of ratio-based approaches to quantitatively assess the early, peak, and terminal phenophases from herbarium specimens and determine phenological changes within and between seasons (Love et al., 2019; Williams et al., 2017). The recent large-scale deployment of the *CrowdCurio* pipeline on Amazon’s Mechanical Turk has demonstrated the power and scale of such fine-grained phenophasing to understand latitudinal variation in phenological responses (Park et al., 2019).

Despite the great promise of crowd-sourcing for phenophase detection, it is still time-consuming and can become cost-prohibitive to process entire collections spanning whole continents. Machine-learning approaches have the potential to open up new opportunities for phenological investigation in the era of Digitization 2.0 (Pearson et al., 2020). Recent efforts (Lorieul et al., 2019) have demonstrated that fully automated machine-learning methods–and deep learning approaches based on convolutional neural networks in particular–can determine the presence of a fruit or flower in a specimen with *>* 90% accuracy. Convolutional neural networks were proven effective at predicting all phenophases of a specimen, based on classification of nine phenological categories. These predictions, estimated from proportions of buds, flowers and fruits, reach an accuracy (true positive rate) *>* 43%, which is equivalent to the capability of human experts (Lorieul et al., 2019). This large-scale automated phenophase estimation, based on an annotation method developed by Pearson (2019), was tested on species belonging to a particularly difficult taxon (*i*.*e*., the Asteraceae family), for which visual analysis of numerous and tiny reproductive structures is known to be visually challenging. This work demonstrated the potential of deep learning technologies to estimate fine-grained phenophases, but further improvements are needed to support ecological investigation of diverse taxa.

Although Pearson (2019) successfully determined reproductive status (*i*.*e*., fertile *vs*. sterile specimens), neither the precise location (*i*.*e*., image segment) nor the number of phenofeatures present on a specimen was quantified (Lorieul et al., 2019). A quantitative machine-learning approach would have the value and impact that *CrowdCurio* has already achieved, but could be scaled-up in speed and cost-effectiveness. A recent proof-of-concept study (Goëau et al., in press) used human-scored data to train and test a model using instance segmentation with Mask R-CNN (Masked Region-based Convolutional Neural Network: He et al., 2017) to locate and count phenological features of *Streptanthus tortuosus* Kellogg (Brassicaceae). This assessment clarified several determinants of model success for identifying and counting phenological features, including: the type of masking applied to human annotations; and the size and type of reproductive features identified (*e*.*g*., flowering buds, flowers, immature and mature fruits). Moreover, the model was more successful identifying and counting flowers than fruits, and was applied only to a single species with relatively little human-scored training data (21 herbarium specimens). The transferability of this model to other, more distantly related species was not examined.

Here, we leverage extensive data gathered using our crowd-sourcing platform *CrowdCurio* to develop and evaluate an instance segmentation approach using Mask R-CNN to train and test a model to identify and count phenological features of a larger number of species. Specifically, we investigated digitized specimens from six common spring-flowering herbs of the eastern United States: *Anemone canadensis, A. hepatica, A. quinquefolia, Trillium erectum, T. grandiflorum*, and *T. undulatum*. As with any feature-detection model, accurate human-collected data are required to train, test, and refine these models. We thus gathered phenological data from these species using *CrowdCurio* to provide expert annotation data of buds, flowers, and fruits to train and test our models. Phenological data previously collected by non-expert citizen scientists was used to further evaluate the performance of these models (Park et al., 2019). Our goals were to: (1) determine how reliably we could localize and count these features; (2) determine the accuracy in automated scoring of different phenological features; and (iii) assess the transferability of models trained on one species to other, distantly related ones.

## 3 Materials and Methods

### 3.1 Dataset

Our experiments are based on a subset of the data used in Park et al. (2018, 2019) comprising six species in two genera of common spring-flowering herbs, *Anemone* and *Trillium*. This subset includes 3073 specimens of: *Anemone canadensis* (*N* = 108), *A. hepatica* (*N* = 524), *A. quinquefolia* (*N* = 686), *Trillium erectum* (*N* = 862), *T. grandiflorum* (*N* = 226), and *T. undulatum* (*N* = 667). Each specimen (herbarium sheet) was previously examined using the *CrowdCurio–Thoreau’s Field Notes* platform by, on average, three citizen-scientists. For the purposes of this study, these specimens were additionally scored by expert botanists to provide the most accurate training and testing data possible. Annotators added markers in the center of each visible reproductive structure (bud, flower, or fruit), and determined its type, number, and spatial location. For our experiments, we randomly split this dataset into two parts: one (*N* = 2457) for training the deep-learning models and one for testing them (*i*.*e*., for evaluating their predictive performance; *N* = 615).

Apart from the comparative experiment described in *§*4.5, only the annotations of experts were used to train and test the deep-learning models. We also only used the annotations of one of the experts for each specimen (selected in a pre-defined order). The final dataset contains 7909 reproductive structures (6321 in the training set and 1588 in the test set) with the following imbalanced distribution: 492 buds (6.2%), 6119 flowers (77.4%), and 1298 fruits (16.4%). Fruits were counted without any knowledge of seeds.

### 3.2 Deep-learning framework

Several deep-learning methods have been developed in recent years to count objects in images. One family of methods can be qualified as density-oriented methods (Zhang et al., 2015; Wang et al., 2015; Boominathan et al., 2016). They are usually based on U-Net architectures (Ronneberger et al., 2015) that are trained on annotations of object centers (indicated by dots) and predict density maps that are integrated to obtain counts. U-Net-based methods were developed originally for counting crowds and have been extended recently to counting cells (Falk et al., 2019) and animals (Arteta et al., 2016). The drawback of these methods is that they are better suited for cases where the density of objects in the image is high. This is not true in our case; the examined herbarium specimens averaged *<* 3 objects per specimen, even fewer if we consider buds, flowers, and fruits separately.

Another deep-learning method is “direct counting” (*a*.*k*.*a*. “glancing”), which trains the model with the true count on the global image (*e*.*g*., Seguí et al., 2015). The main drawback of direct counting is that it cannot predict a count value that has no representative image in the training set. That is, the network is not really counting but only inferring the counts from the global content of the image. In preliminary experiments (not reported here), we found that direct-count methods tended to systematically under-estimate the true counts and have an unacceptably high variance.

The alternative method that we used in this study is to equate counting with object-detection; the counts of the object of interest is then equal to the sum up the number of detected objects. To detect buds, flowers, and fruits, we used Mask R-CNN, which is among the best-performing methods for instance segmentation tasks in computer vision (He et al., 2017). We used Facebook’s implementation of Mask R-CNN (Massa and Girshick, 2018) using the PyTorch framework (Paszke et al., 2019) with a ResNet-50 architecture (He et al., 2016) as the backbone CNN and the Feature Pyramid Networks (Lin et al., 2017) for instance segmentation. To adapt this architecture to the data in our study (see previous section), we had to address the following methodological issues:

1. **Mask computation**. The training data expected by Mask R-CNN must consist of all the objects of interest visible in the training images, each object being detected individually and associated with a segmented region (encoded in the form of a binary mask). However, the data available for our study did not fully meet these conditions as the objects were detected only by dot markers (roughly in the centre of the reproductive structure). From these dot markers, we generated dodecagons, such as the ones illustrated in Figure 1, which best covered the reproductive structures. To adapt the size of the dodecagons to buds, flowers, and fruits, we manually segmented five of each (selected at random from each genus) and calculated the average radius of the circle enclosing each structure.
2. **Input image size**. Images were resized to 1024 pixels (long edge) *×* 600 pixels (short edge). This guaranteed a sufficient number of pixels for the smallest dodecagons while maintaining a reasonable training time (5–10 hours per model) on a computer comparable to a mid-tier consumer device (*i*.*e*., recent GPUs with *±*12 GB of RAM).
3. **Anchor size**. Anchors are the raw rectangular regions of interest used by Mask R-CNN to select the candidate bounding boxes for mask detection. We designated their size so as to guarantee that all dodecagons had their entire area covered.

Figure 2 illustrates four example detections using Mask R-CNN: one with a perfectly predicted count, and three with over- or under-estimated counts. For each example, we show (a part of) the original image, the ground-truth masks (computed from expert botanist input), and the automated detections computed by the deep-learning framework.

**Figure 1:**
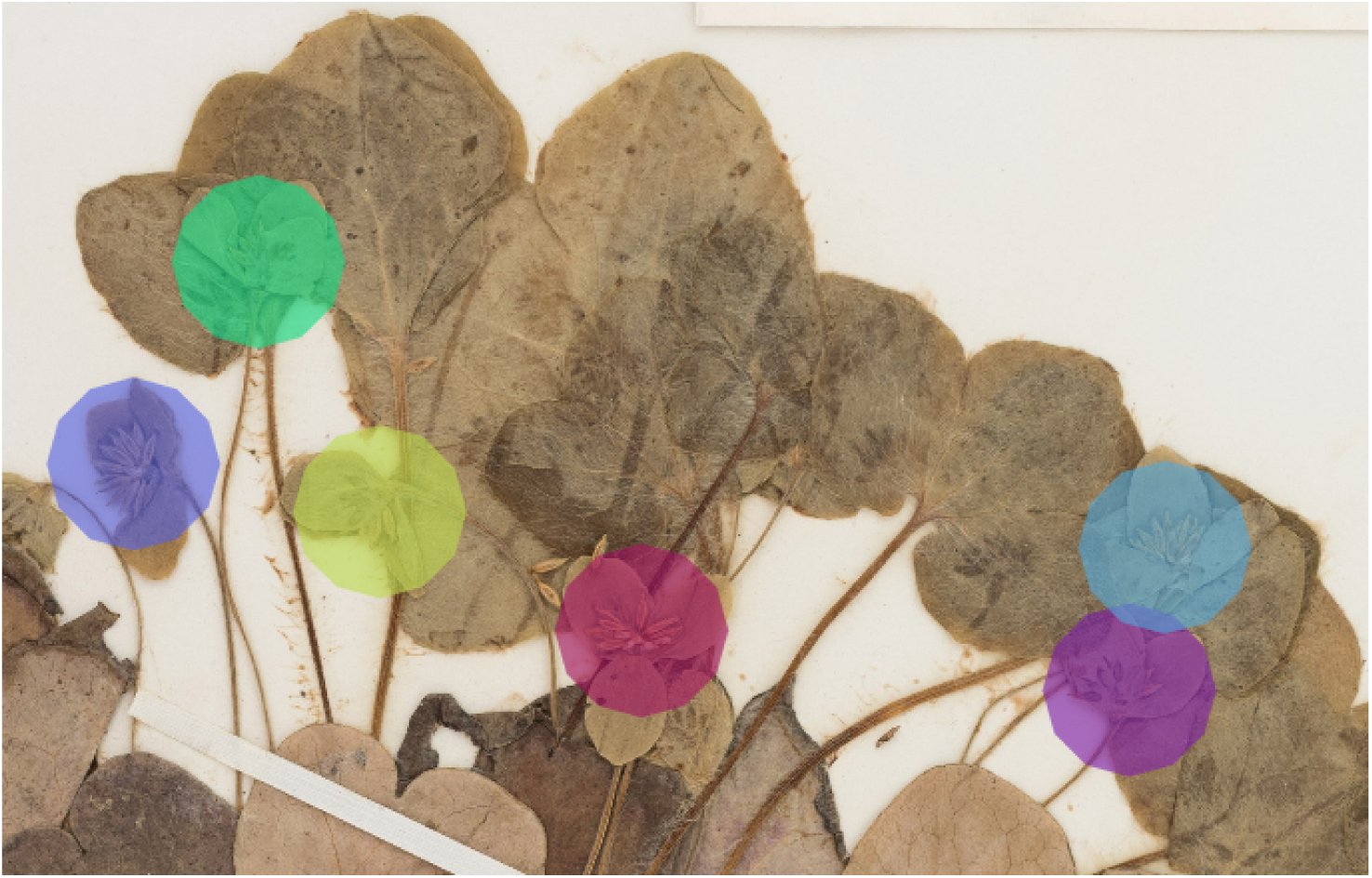
Example of a specimen of the training set containing six reproductive structures (flowers) marked by dodecagons.

**Figure 2:**
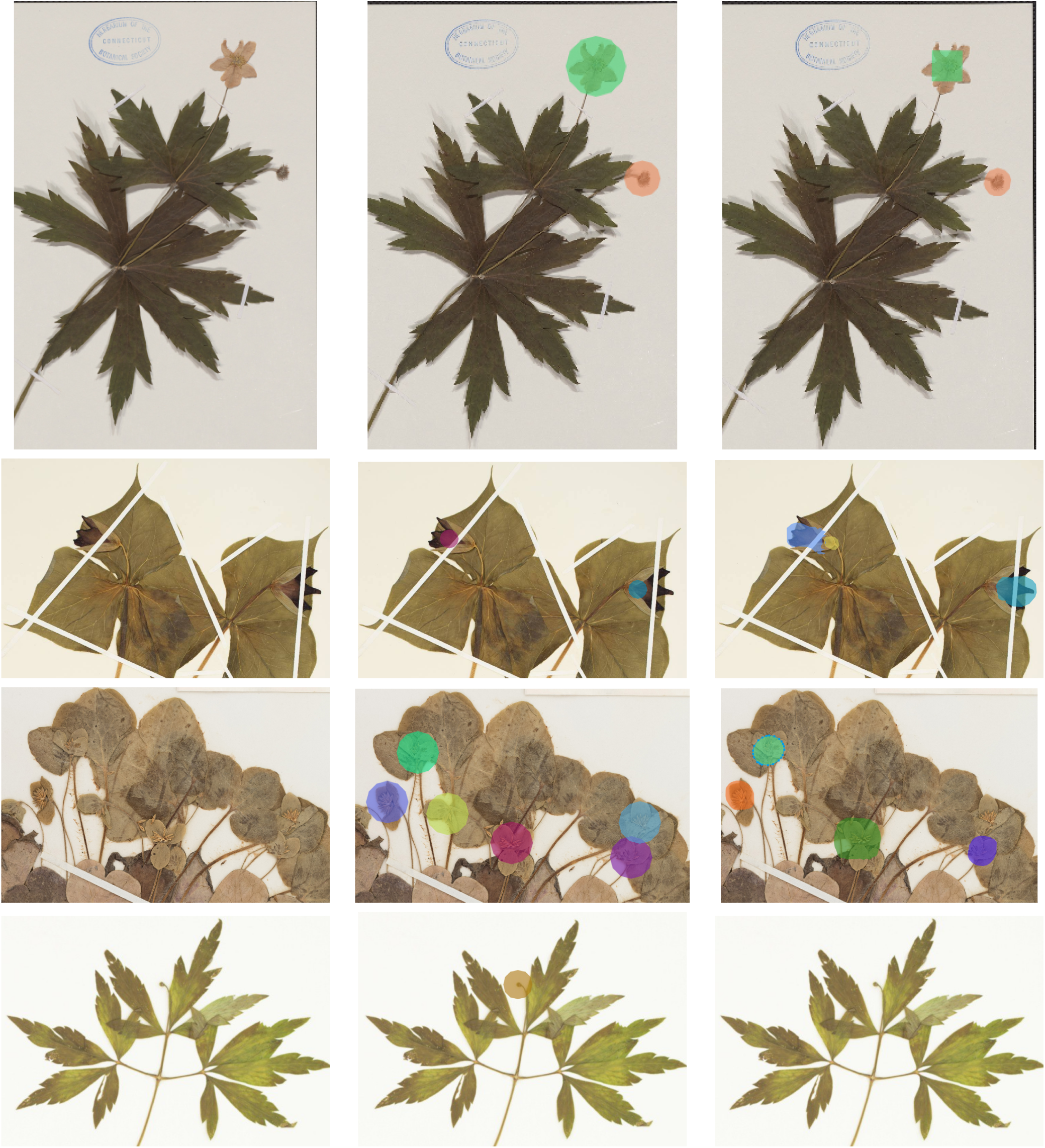
Examples of detection (colors do not have a particular meaning) - **Left Column**: original image; **Center Column**: ground-truth markers; **Right Column**: automatically detected masks. The first row corresponds to a typical case with a perfect count. The second row corresponds to a case of over-estimated counts (one of the flowers was detected as two flowers). The last two rows correspond to under-estimated counts (some structures were missed or aggregated as one).

We then trained a set of models corresponding to three distinct scenarios to be evaluated:

1. **One model per species**. In this scenario, we trained one Mask R-CNN model for each species (*i*.*e*., six models in total) to detect its buds, flowers, and fruits.
2. **One single model for all species**. In this scenario, we trained a single Mask R-CNN for all species and all types of reproductive structures (buds, flowers, fruits).
3. **Cross-species models**. Last, we assessed the transferability of models trained on some species to other ones. We trained three models on only two *Trillium* species: *i*.*e*., one on *T. erectum* and *T. grandiflorum*, one on *T. erectum* and *T. undulatum*, and one on *T. undulatum* and *T. grandiflorum*. Each of these three models were then tested on the *Trillium* species not included in the training set.

### 3.3 Evaluation metrics and statistics

We evaluated the accuracy of the models in four ways:

1. **Counting error**. The counting error *e*_*i,k*_ for a specimen *i* and a given type of reproductive structure *k* ∈ {*bud, flower, fruit*} was defined as the difference between the true count and the predicted count:

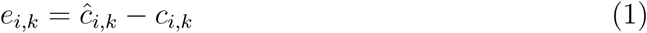

where *c*_*i,k*_ is the true count of reproductive structures of type *k* in specimen *i* and *ĉ*_*i,k*_ is the predicted count. Note that the counting error can be positive or negative. A detailed description of the distribution of the counting error is provided using letter-value plots (Heike et al., 2017), which provide a more comprehensive view of the statistics through a larger number of quantiles.
2. **Mean Absolute Error (MAE)**. The MAE measures the overall error by averaging the absolute value of the counting error of each specimen and each type of reproductive structure:

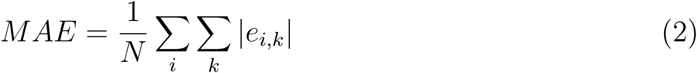
3. **Coefficient of determination (***R*^2^). This statistic measures the amount of variance explained or accounted by the model:

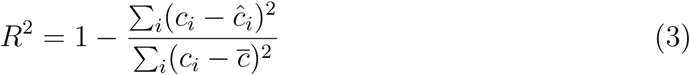

where *i* indexes the observations and ranges from 1 to the total number of observations, *c*_*i*_ is the observed count, *ĉ*_*i*_ is the predicted count, and 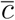 is the mean of the observed counts.
4. **Predicted counts box-plots**. A detailed description of the distribution of the predicted counts as a function of the true counts is provided using box-plots indicating median value, quartiles, variability outside quartiles, and outliers.

### 3.4 Machine-learning *vs*. crowd-sourcing

We compared the counts predicted by Mask R-CNN with those obtained when the reproductive structures on herbarium specimens were counted by crowd-sourcers (Park et al., 2019). The comparison was done on the intersection of the test sets of both studies (*i*.*e*, on 544 specimens, equal to 88% of the test set of previous experiments). These 544 specimens were annotated by 483 different annotators using Amazon Mechanical Turk. On average, each specimen was annotated by 2.5 different crowd-sourcers.

## 4 Results

### 4.1 A single model *vs*. species-specific models

The *R*^2^ value for the separate training model for each species and the single model for all species was 0.70 and 0.71, respectively. Thus, the single model for all species provides marginally better results while being simpler to implement and more scalable. As shown in Fig. 3, the main problem of single species training models is that they tend to over-predict the number of reproductive structures (number of positive errors *>* than number of negative errors; Fig. 3). The extreme outlier in Fig. 3 with a very high negative error resulted from a species being assessed by the model that had been misidentified in the collection.

**Figure 3:**
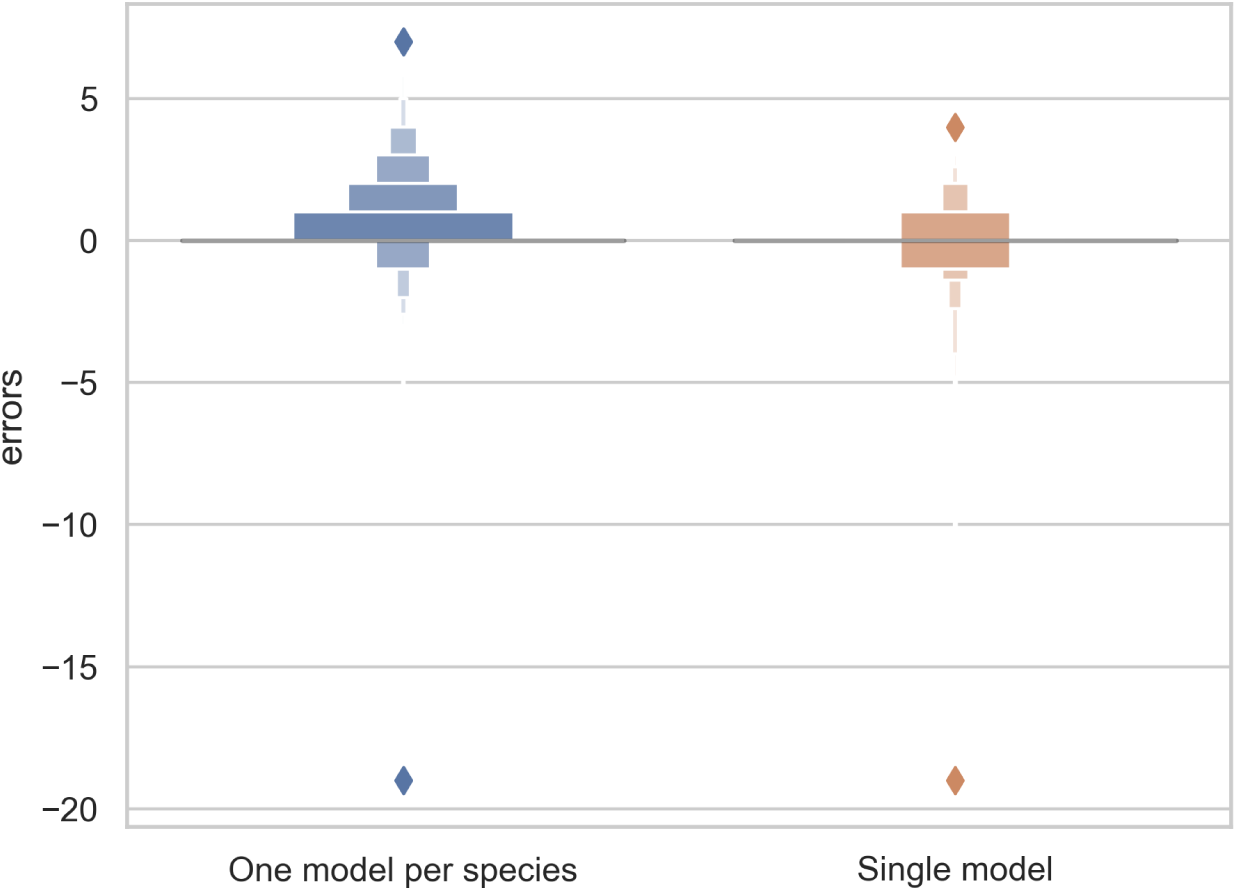
Letter-value plot of the counting error for the two training strategies: one model per species *vs*. one single model for all species.

The predictions of the single species training models were very accurate for ≤ 3 reproductive structures, whereas the single model for all species had high accuracy when ≤ 4 reproductive structures were present (Fig. 4). The variance of the predicted counts was higher for specimens with more reproductive structures but the median predicted count equalled the actual count for ≤ 7 reproductive structures and the counting error (interquartile distance) was usually *<* 1 structure. Specimens with *>* 8 reproductive structures had larger errors but only accounted for 4.2% of the specimens examined.

**Figure 4:**
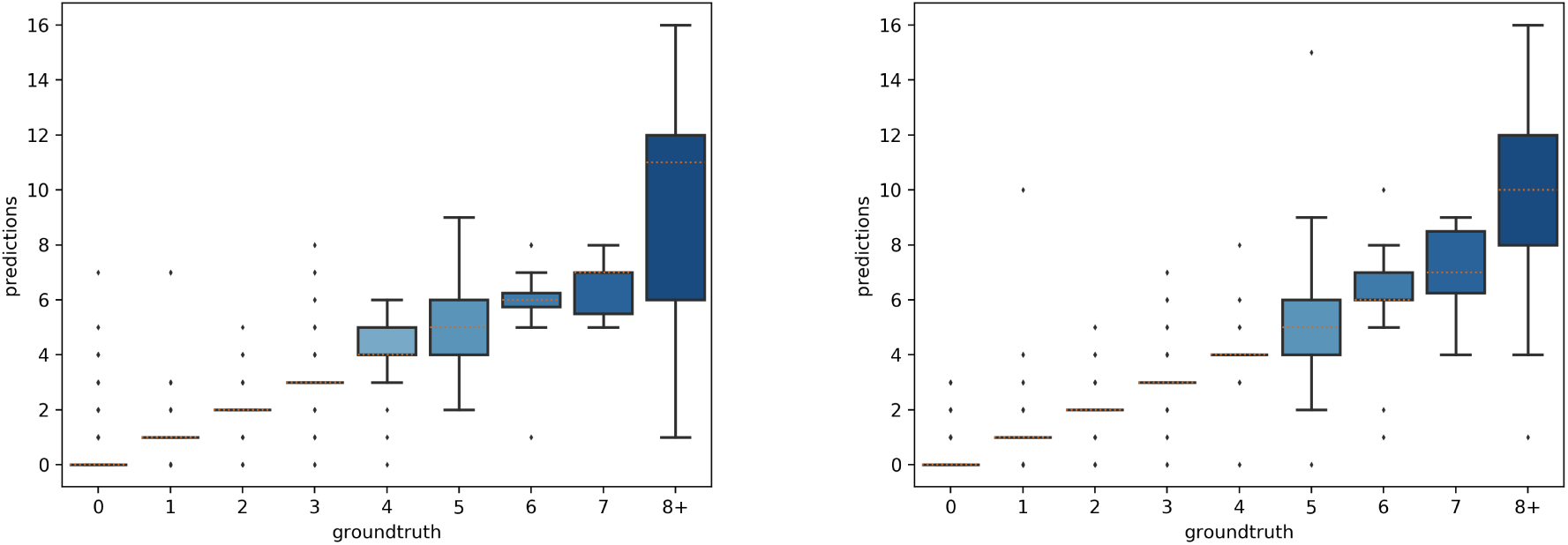
Box-plots of the predicted *vs*. expected counts for the two training strategies: **(Left)** separate training models for each species, **(Right)** single training model for all species.

### 4.2 Distinguishing reproductive structures

#### 4.2.1 Counting results

The overall numbers of detected reproductive structures and their relative proportions were very close to their actual values (Table 1). The Mean Absolute Error (MAE) was also quite low for all types of reproductive structures, but this is due in large part to the fact that the median number of structures per phase and specimen is low. The median number of fruits and buds, in particular, is much lower than the median number of flowers. The *R*^2^ values (Table 1) and the box plots of the predicted counts (Fig. 6) provide a more relevant comparison of the predictive performance for each type of structure. Flowers are the best detected structures (*R*^2^ = 0.76), followed by fruits (*R*^2^ = 0.33) and buds (*R*^2^ = 0.12). The lower performance for buds is due to several factors: (i) the lower number of samples in the training set–90.25% of specimens had no buds and 98.05% had *<* three buds, (ii) their smaller size and (iii), their visual appearance that is less distinctive than flowers or fruits. Fruits are affected by the same factors but to a lesser extent.

**Table 1:** Predicted and true counts (percent of specimens in parentheses) of buds, flowers, and fruits for all specimens pooled.

|  | Buds | Flowers | Fruits | All |
| --- | --- | --- | --- | --- |
| True number of structures | **107** (6.7) | **1241** (78.1) | **240** (15.1) | 1588 |
| Predicted number of structures | **109** (6.1) | **1431** (80.0) | **248** (13.9) | 1788 |
| MAE | **0.20** | **0.51** | **0.27** | **0.33** |
| $R^2$ | **0.12** | **0.76** | **0.33** | **0.71** |

**Figure 5:**
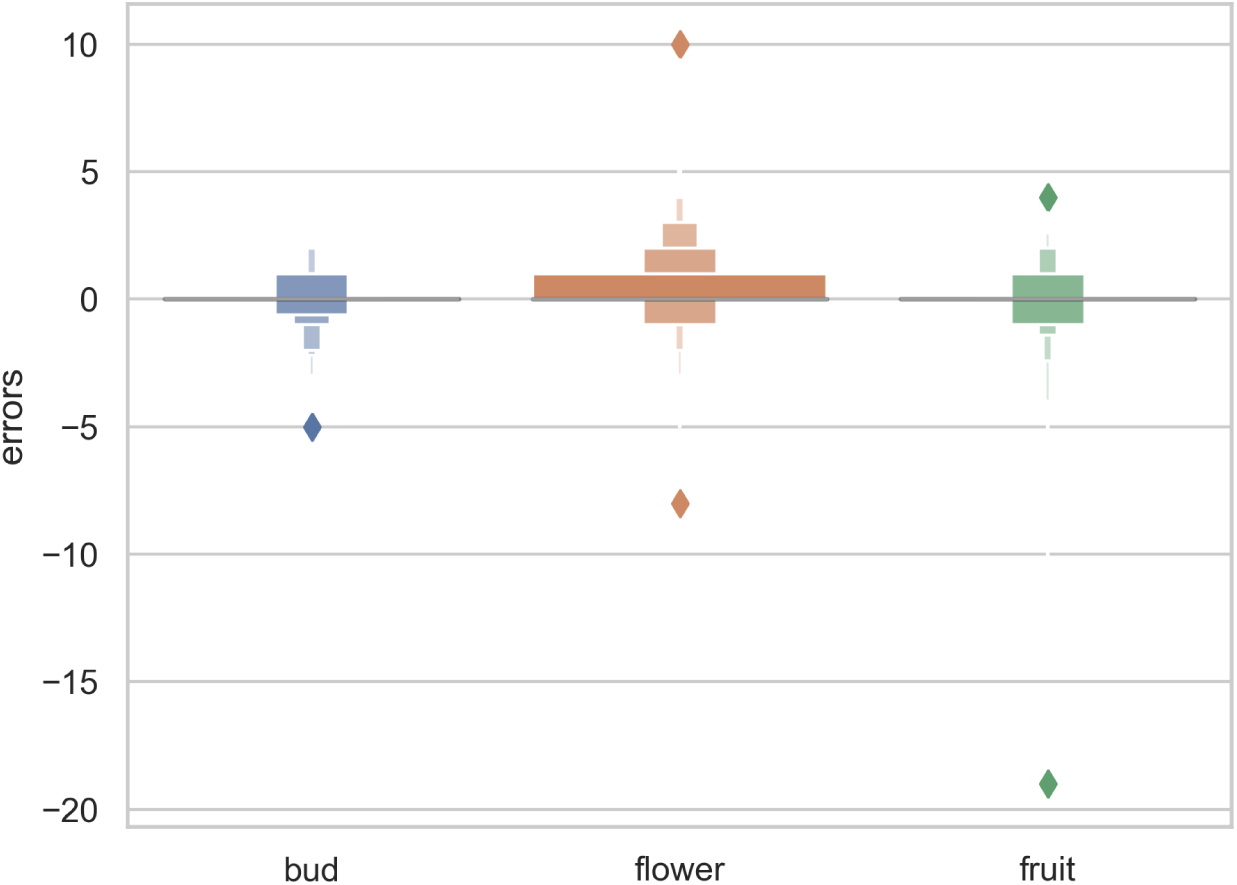
Letter-value plot of the counting error for each type of reproductive structure.

**Figure 6:**
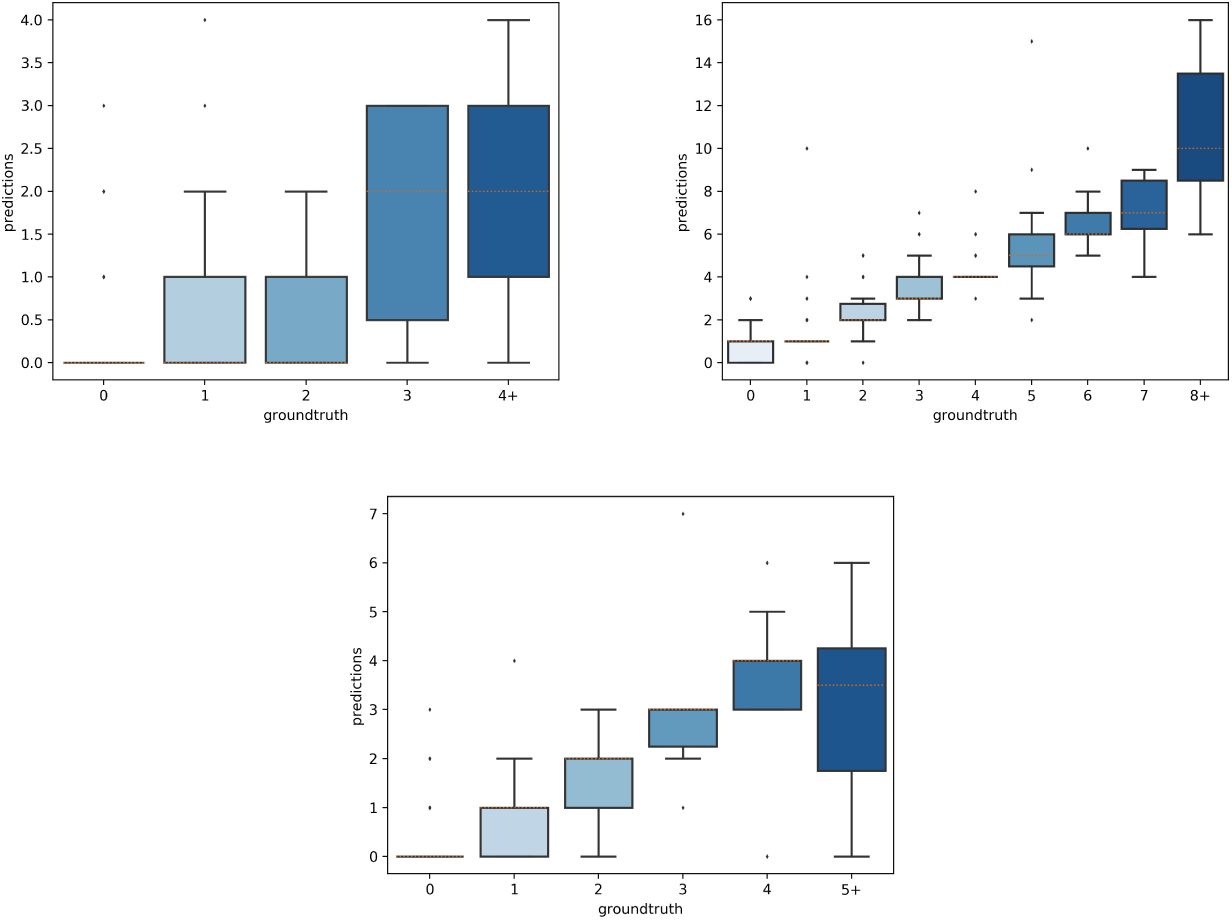
Box-plots of the predicted *vs*. expected counts for each type of reproductive structure. From left to right: buds, flowers, fruits.

#### 4.2.2 Occurrence and dominance of reproductive structures

Although the model was not developed or trained to directly detect presence or absence of each reproductive structure, we were able to extrapolate the presence of each feature and which feature was most frequent on a specimen (Table 2). The detection accuracy of buds, flowers, and fruits was *>* 87% and the accuracy of determining relative abundance of a certain organ category (*e*.*g*., number of flowers *>* number buds or fruits) was *>* 90% (Table 2). Confidence in this strong result should be tempered by the actual frequency of occurrence and dominance. Observed relative presences of buds, flowers, and fruits, and dominance of fruits *vs*. flowers all are quite disparate. Error rates (false negatives and positives) for these all are non-zero, but are lower in all presence and dominance categories (Table 2).

**Table 2:** Accuracy of detection and relative dominance of buds, flowers, and fruits (data pooled for all species). Values are percentages.

| | Buds | Flowers | Fruits | Flowers≥Buds | Fruits≥Flowers |
| --- | --- | --- | --- | --- | --- |
| **Observed** | 9.75 | 82.92 | 20.00 | 96.09 | 21.13 |
| True positives (correctly detected) | 51.66 | 97.25 | 78.86 | 98.98 | 76.15 |
| True negatives (correctly undetected) | 91.89 | 49.52 | 89.83 | 8.33 | 95.65 |
| False positives | 8.10 | 50.47 | 10.16 | 91.66 | 3.71 |
| False negatives | 48.33 | 2.74 | 21.13 | 1.01 | 23.84 |
| **Overall Accuracy** | **87.97** | **89.11** | **87.64** | **95.44** | **92.03** |

### 4.3 Species-specific models

Overall, the reproductive structures were detected more accurately for *Trillium* species than *Anemone* species (Figs. 7 and 8). At the species-specific level, the *R*^2^ score was lowest for *A. canadensis* (0.01) which is the species with the least number of training samples (108 specimens). The *R*^2^ score was better for the other species and increased with the number of training samples: *R*^2^ = 0.51 for *T. grandiflorum, R*^2^ = 0.64 for *A. hepatica, R*^2^ = 0.76 for *T. undulatum, R*^2^ = 0.85 for *A. quinquefolia* and *R*^2^ = 0.89 for *T. erectum*. Counting errors rarely exceeded *±*2, and the few strong outliers corresponded to very difficult cases or annotation errors. The median value of predicted counts was correct in almost all cases (Fig. 7); exceptions were for *T. grandiflorum* specimens with four structures and *A. hepatica* with seven, both corresponding to instances involving a small number of specimens with large numbers of reproductive structures.

**Figure 7:**
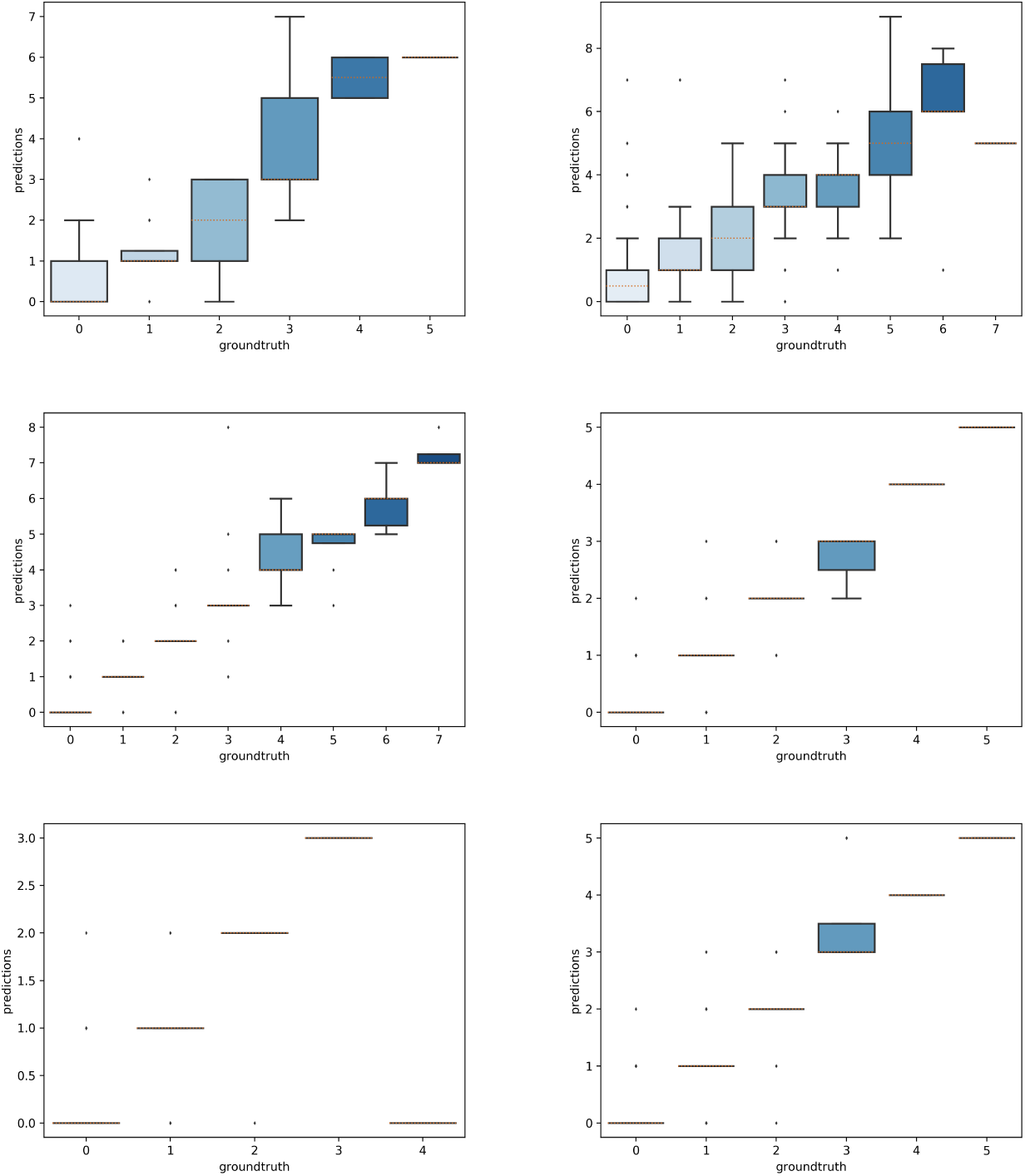
Boxplot of the predicted counts vs. expected counts for each species. **(A)**: *Anemone canadensis*; **(B)**: *A. hepatica*; **(C)**: *A. quinquefolia*; **(D)**: *Trillium erectum*; **(E)**: *T. grandiflorum*; **(F)**: *T. undulatum*.

**Figure 8:**
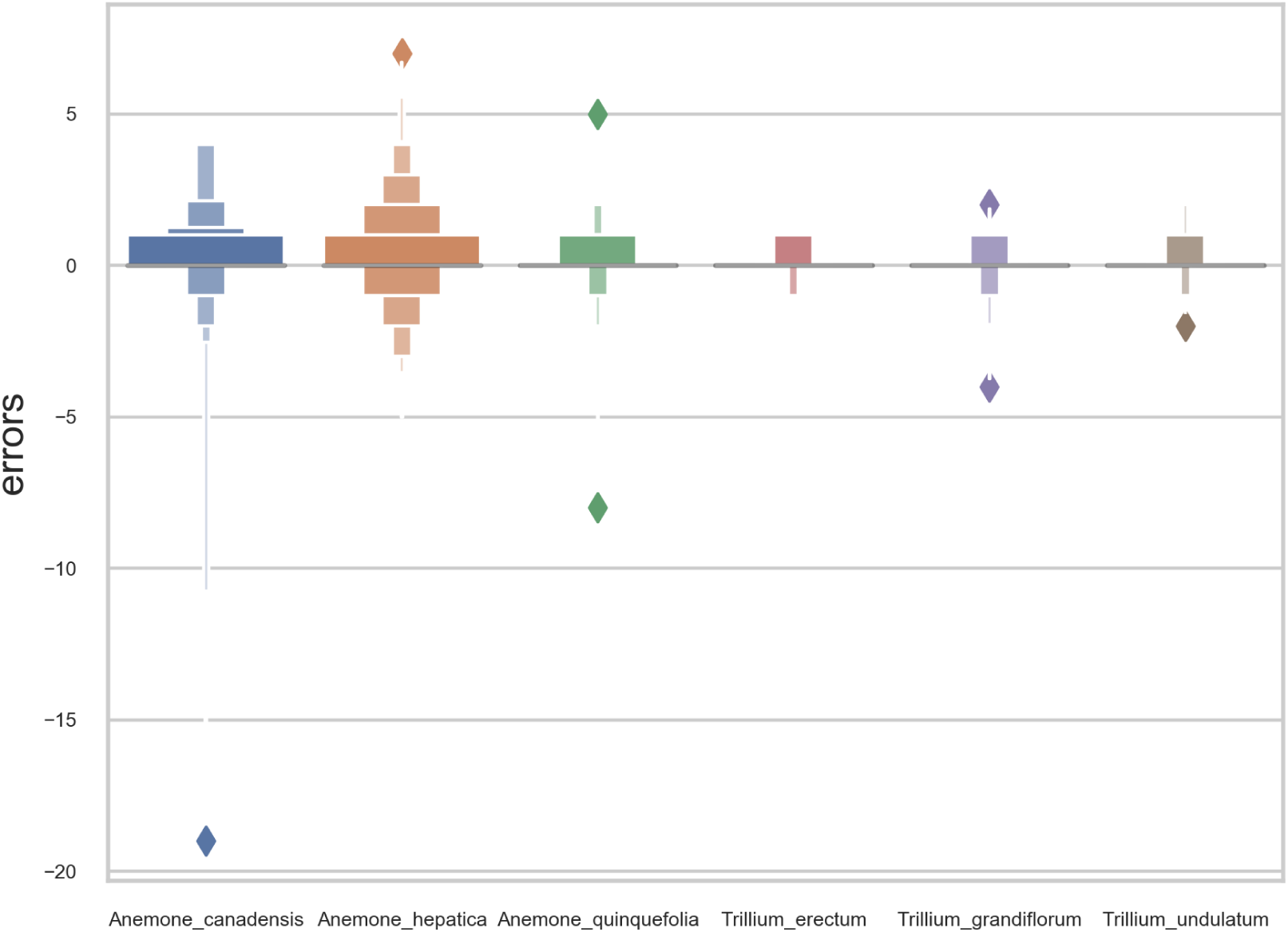
Letter-value plot of the counting error for each species.

### 4.4 Model transferability

The aim of this experiment was to assess whether reproductive structures on one species could be estimated using a model trained on a different, related species. Unsurprisingly, estimation was less accurate when the target species was not represented in the training set (Figs. 9–11). However, it is still possible to count the reproductive structures of a target species based on a model trained on different species of the same genus (*i*.*e*., without any specimen of the target species in the training data). The *R*^2^ score was higher for *T. erectum* (*R*^2^=0.72; Fig. 9) and *T. undulatum* (*R*^2^=0.66; Fig. 10), which are morphologically more similar to one another than either is to *T. grandiflorum* (*R*^2^=0.02; Fig. 11). Figures only show the results for *Trillium* but similar conclusions were obtained for *Anemone* (*R*_2_ scores respectively equal to 0.75 for *A. quinquefolia*, 0.39 for *A. hepatica* and -0.39 for *A. canadensis*).

**Figure 9:**
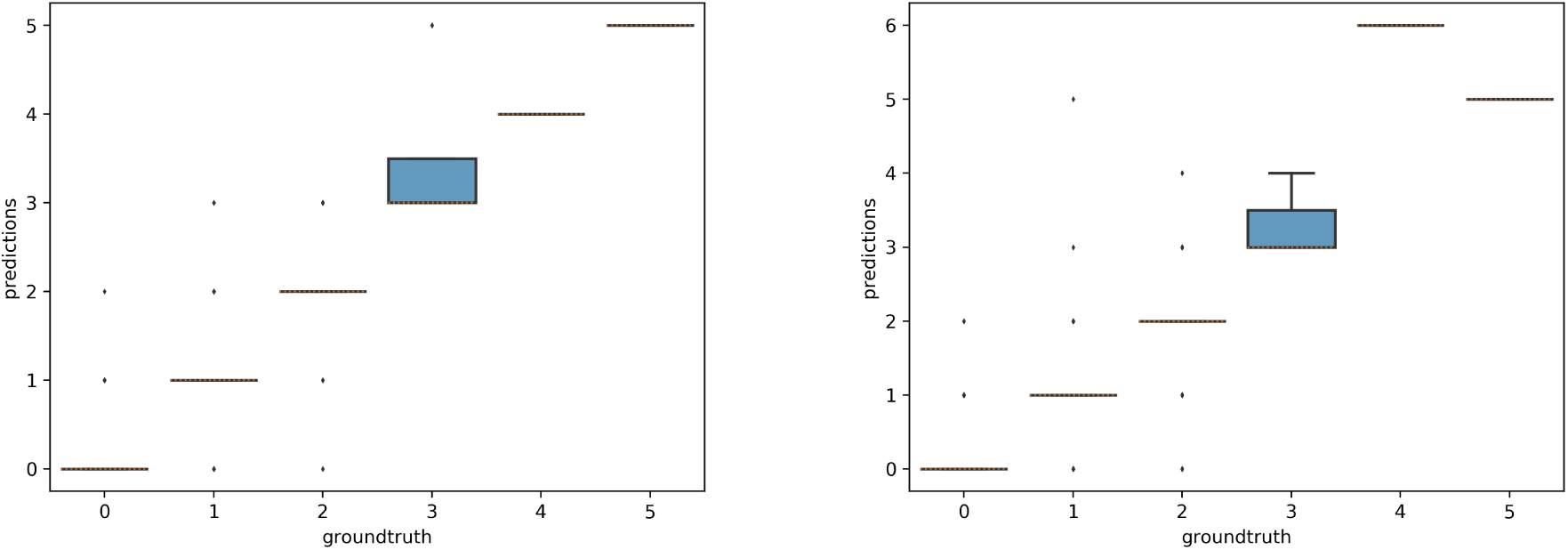
Box-plots of the predicted counts *vs*. expected counts for *Trillium erectum*. Left: Model trained on *T. erectum* data; Right: model trained on *T. undulatum* and *T. grandiflorum*.

**Figure 10:**
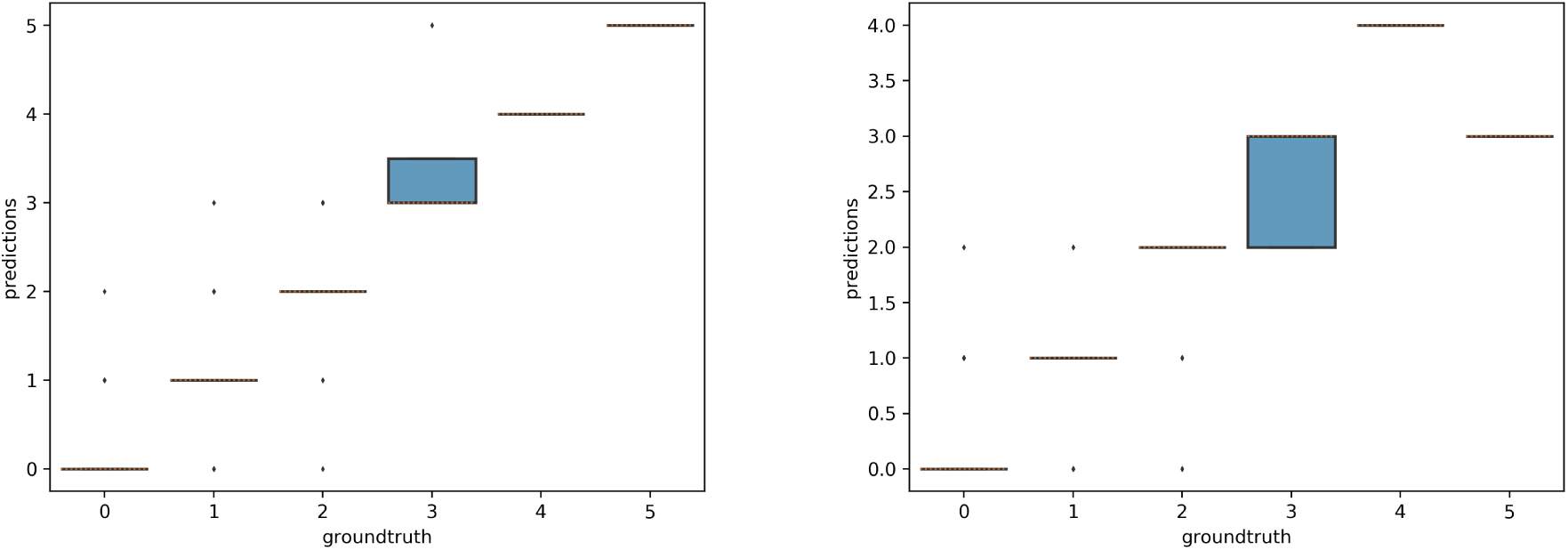
Box-plots of predicted counts *vs*. observed counts for *Trillium undulatum*. Left: Model trained on *T. undulatum* data; Right: model trained on *T. erectu*m and *T. grandiflorum*.

**Figure 11:**
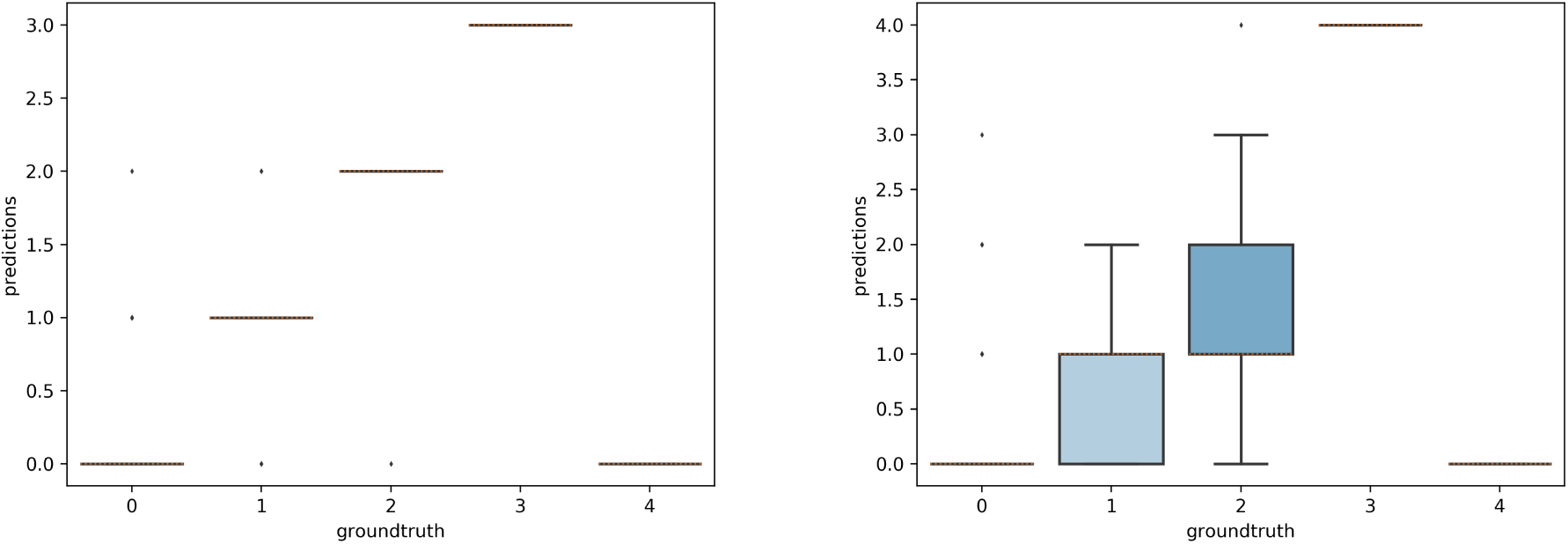
Box-plots of predicted counts *vs*. expected counts for *Trillium grandiflorum*. Left: Model trained on *T. grandiflorum* data; Right: model trained on *T. erectum* and *T. undulatum*

### 4.5 Machine-learning *vs*. crowd-sourcing

On average, the deep learning model had a significantly lower (*P <* 0.001) MAE and better *R*^2^ score than any individual crowd-sourcer, but still an order of magnitude larger than the MAE of botanical experts (Table 3 and 4). Interestingly, we can observe that crowd-sourcers have a much harder time detecting buds than the Mask R-CNN model. The MAE obtained by averaging the counts of the different crowd-sourcers was only marginally higher than the MAE from Mask R-CNN (*P* = 0.3). Note that a counts averaging strategy could also be used for the deep learning approach, *i*.*e*., by averaging the scoring of several deep learning models. This technique is referred to as an *ensemble* of models in the machine learning community and is known to bring very significant improvements. The most simple yet very efficient method to build an ensemble is to train several times the same model but with a different random initialization of the parameters. Such strategy could be implemented in future work.

**Table 3:** Comparison of the counting error resulting from crowd-sourcing, deep learning and expert annotation – performance is measured by the Mean Absolute Error (MAE).

|  | Buds | Flowers | Fruits | All |
| --- | --- | --- | --- | --- |
| Experts | 0.009 | 0.027 | 0.073 | 0.036 |
| Crowd-sourcing (isolated annotator) | 0.526 | 0.487 | 0.314 | 0.442 |
| Crowd-sourcing (average over all annotators) | 0.418 | 0.405 | 0.243 | 0.355 |
| Deep learning (model trained on all species) | **0.201** | **0.507** | **0.266** | **0.325** |

**Table 4:** Comparison of the counting error resulting from crowd-sourcing, deep learning and expert annotation – performance is measured by *R*^2^ score.

|  | Buds | Flowers | Fruits | All |
| --- | --- | --- | --- | --- |
| Experts | 0.989 | 0.996 | 0.961 | 0.990 |
| Crowd-sourcing (isolated annotator) | -2.969 | 0.758 | 0.306 | 0.555 |
| Crowd-sourcing (average over all annotators) | -1.527 | 0.828 | 0.401 | 0.686 |
| Deep learning (model trained on all species) | **0.141** | **0.750** | **0.329** | **0.707** |

## 5 Discussion

Mask R-CNN models trained with human-annotated trait data were efficient and produced robust results. Our models worked well for both identifying and counting phenological features, but accuracy differed for buds, flowers, and fruits. Automated counts using Mask R-CNN models were more accurate than counts made by crowd-sourcers but not those of botanical experts. Finally, the Mask R-CNN model could be transferred to other species after being trained with data from reasonably close phylogenetic relatives, with relatively small impacts on counting accuracy.

### Point masking with minor modification is efficient and produces robust results

Recent efforts by Goëau et al. (in press) to segment and count reproductive structures used training data collected by botanical experts from 21 herbarium specimens of a single species (*Streptanthus tortuosus*). In our work, we applied Mask R-CNN to segment and count reproductive structures of six species, belonging to two different genera; accurate training data were derived from both botanical experts and crowd-sourcers using the *CrowdCurio* interface (Willis et al., 2017). Although Goëau et al. (in press) found that training data from point masks, like those generated from *CrowdCurio*, produced less accurate results than those derived from fully masked training data, obtaining the latter is time intensive and difficult to scale to large numbers of specimens. Whereas Goëau et al. (in press) produced three type of training data, “point masks” (produced from a 3 *×* 3-pixel box around a manual point marker); (ii) “partial masks” (extensions of point masks to include partial segmentation using the Otsu segmentation method (Otsu, 1979); and (iii) manually produced “full masks” of each reproductive structure, we only used modified partial masks (derived from point markers) with Mask R-CNN. These modified partial masks were scaled to the size of reproductive structures for each species and yielded high accuracy and efficiency for phenophase detection and counting. The scaling of our modified partial masks combined with the approximately circular shapes of the reproductive structures we studied likely led to the success of our approach. Our two-step workflow integrating expert-scored and crowd-sourced citizen science data with automated machine-learning models also is less time-intensive and more scalable than a workflow requiring detailed polygon masks of structures for training.

### Feature detection and counting accuracy is high across all phenological features

Lorieul et al. (2019) were the first to apply machine-learning to detect phenophases and developed a presence-absence model that could identify reproductive specimens with ≈ 96% accuracy. Their model was less accurate in detecting flowers or fruits (≈ 85% and ≈ 80% accuracy, respectively), and they did not consider buds. In contrast, we used Mask R-CNN to accurately identify the presence of each of the three reproductive stages (buds, flowers, or fruits) with ≥ 87% accuracy (Table 2). Moreover, a single globally-trained model was more efficient and had greater accuracy than multiple species-specific models (Figs. 7 and 8). This points towards the possibility of developing a more streamlined workflow to accurately score phenophases of many different species simultaneously.

We also successfully estimated the relative abundance of each reproductive structure on a specimen with ≥ 90% accuracy (Table 2). Herbarium specimens can vary greatly in phenological state. Because different reproductive organs can co-exist at various times through plant development (and may not all be represented simultaneously on herbarium sheets), simply quantifying presence or absence of phenological structures limits inference about phenological state. In this regard, the Mask R-CNN model performed better on *Trillium*—with its large flowers and fruits, generally borne singly, and suspended on an elongate stalk—than on *Anemone*—with its small clusters of flowers on shorter stalks that are often pressed against a background of clustered leaves. The combination of smaller flowers, more complex morphology, and background “noise” on *Anemone* specimens (*e*.*g*., overlapping structures) likely made both model training and phenophase detection more prone to error. This result supports the recent hypotheses that successful application of machine-learning to phenophase assessment will be dependent on species-specific morphological details (Goëau et al., in press). Along these lines, plant morphological trait databases could help facilitate the identification of suitable taxa to be analysed with machine-learning methods.

Precise quantification of different reproductive structures, as demonstrated here, allows the determination of finer-scale phenophases (*e*.*g*., early flowering, peak flowering, peak fruiting). For this exercise, the lowest mean absolute error (MAE) was for bud counts, most likely due to the morphological consistency of buds and their rarity on specimens (Table 1). In contrast, MAE for counting flowers was significantly worse than for buds or fruits. We attribute this result to the greater number of flowers, ontogenetic variability in floral morphology, and variation in appearance of dried, pressed specimens.

Variation in appearance of reproductive features among dried and pressed specimens of a single species also could add complexity to automated detection of phenological features and merits further investigation. Perhaps more consequentially, large variation in the number of reproductive organs resulted in unbalanced datasets (Table 1). Numerous data augmentation approaches can be implemented to improve comparisons and model selection for such data sets (*e*.*g*., Tyagi and Mittal, 2020), but these approaches have been used more frequently in classification or semantic segmentation (Chan et al., 2019) than in instance segmentation approaches such as we used here. Developing data augmentation approaches for instance segmentation would be a useful direction for future research. But even if collectors collect more flowering than non-flowering specimens, estimating the quantity of buds, flowers and fruits on any specimen is more informative than recording only their presence or absence.

### Botanical experts perform better than the model

When considered in aggregate, the MAE for segmenting and counting all three phenophases using Mask R-CNN was lower than that of crowd-sourcers but still an order of magnitude higher than that of botanical experts (Tables 2, 3). This result reinforces the suggestion that abundant and reliable expert data are essential for properly training and testing machine learning models (Brodrick et al., 2019). Additionally, it was evident in some cases that the precise detection of the phenological feature was quite inaccurate (Figure 2).

### Machines can apply learning from one species to another, but success is variable

For the first time to our knowledge, we have demonstrated that training data from related taxa can be used to detect and count phenological features of a species not represented in the training set (Figs. 9–11). We limit our discussion of transferability here to species of *Trillium* owing to the ease of detecting and counting phenological features in this genus. Though in some cases species-specific models were highly transferable, model transferability varied greatly. For example, training on *T. undulatum* and testing on *T. erectum* (and vice-versa) was more accurate than when Mask R-CNN models trained with data from either of these species was applied to *T. grandiflorum. Trillium undulatum* and *T. erectum* are more similar morphologically than either is to *T. grandiflorum*, suggesting that morphological similarity may be a better guide for transferability success than phylogenetic relatedness (see Farmer and Schilling, 2002, for phylogenetic relationships of *Trillium*). This conclusion implies that transferability may be particularly challenging for clades that exhibit high morphological diversity and disparity among close relatives. The relationship between phylogenetic relatedness, morphological diversity, and model transferability should be investigated in future studies. The assessment of the sizes of the reproductive structures that could be captured by this type of approach should also be analysed, to facilitate transferability.

### Future directions

The presence of reproductive structures has been determined only infrequently during large-scale digitization and transcription efforts by the natural-history museums that generate this content. However, interest is growing rapidly in using herbarium specimens for investigating historical changes in phenology and other ecological traits and processes. Our results have demonstrated success in automating the collection of large amounts of ecologically-relevant data from herbarium specimens. Together with controlled vocabularies and ontologies that are being developed to standardize these efforts (Yost et al., 2018), our two-stage workflow has promise for automating and harvesting phenological data from images in large virtual herbaria. In the long term, we would like to use the *CrowdCurio* workflow to generate reliable human-annotated data to further refine automated models for detecting phenological responses to climatic change from herbarium specimens across diverse clades and geographies. Finally, our results documenting transferability of machine-learning models from one species to another are preliminary, but promising. Although our universal model trained on all taxa performed better than our individual, species-specific models, there may be better ways to guide these efforts. For example, a hierarchy of individual models could yield more accurate results. These hierarchies might be phylogenetically organized (*e*.*g*., taxonomically by order, family, genus), leveraging information about shared morphologies common to related taxa and further governed by a set of rules that parse new specimens for phenophase detection based on their known taxonomic affinities (*e*.*g*., by genera). Similar approaches are already being applied today by corporations like Tesla Motors. Their automated driving suite uses different models for vehicle path prediction versus vehicle detection (Karpathy et al., 2014; Tesla, 2019).

## Author Contributions

CCD conceived the idea for the study; CCD, DT, IB, and DSP ran a pilot feasibility study to motivate the current project; DSP and GML generated, organized, and assembled expert and non-expert crowd-sourced data to train the Mask R-CNN model; JX re-coded *CrowdCurio* for these experiments; JC, AJ, and PB conducted the analyses; CCD, AJ, PB, JC, DSP, and AME interpreted the results; CCD wrote the first draft of the Abstract, Introduction, and Discussion; JC, AJ, and PB wrote the first draft of the Methods and Results; all co-authors revised and edited the final draft.

## Acknowledgments

The authors are grateful to Inria Sophia Antipolis - Méditerranée “NEF” computation platform for providing resources and support. The authors acknowledge iDigBio’s Phenology and Machine Learning Workshop (1/2019), which helped to stimulate this collaboration. The authors are grateful for the efforts of citizen scientists that helped generate data and the many collectors and curators of plant specimens that have made this research possible.

## Supplemental Data

### Data Availability Statement

The datasets generated and analyzed for this study can be found in the Harvard Forest Data Archive https://harvardforest.fas.harvard.edu/data-archive, dataset HF-3xx and the Environmental Data Initiative doi:https:/dx.doi.org/doi-to-come.

